# On the Relation of Gene Essentiality to Intron Structure: A Computational and Deep Learning Approach

**DOI:** 10.1101/2020.03.31.019125

**Authors:** Ethan Schonfeld, Edward Vendrow, Joshua Vendrow, Elan Schonfeld

**Author notes:** **Corresponding Author** Correspondence to Ethan Schonfeld.

## Abstract

Identification and study of human-essential genes has become of practical importance with the realization that disruption or loss of nearby essential genes can introduce latent-vulnerabilities to cancer cells. Essential genes have been studied by copy-number-variants and deletion events, which are associated with introns. The premise of our work is that introns of essential genes have characteristic properties that are distinct from the introns of nonessential genes. We provide support for the existence of characteristic properties by training a deep learning model on introns of essential and nonessential genes and demonstrated that introns alone can be used to classify essential and nonessential genes with high accuracy (AUC of 0.846). We further demonstrated that the accuracy of the same deep-learning model limited to first introns will perform at an increased level, thereby demonstrating the critical importance of introns and particularly first introns in gene essentiality. Using a computational approach, we identified several novel properties of introns of essential genes, finding that their structure protects against deletion and intron-loss events, and that these traits are especially centered on the first intron. We showed that GC density is increased in the first introns of essential genes, allowing for increased enhancer activity, protection against deletions, and improved splice-site recognition. Furthermore, we found that first introns of essential genes are of remarkably smaller size than their nonessential counterparts, and to protect against common 3’ end deletion events, essential genes carry an increased number of (smaller) introns. To demonstrate the importance of the seven features we identified, we trained a feature–based model using only information from these features and achieved high accuracy (AUC of 0.787).

## Introduction

Essential genes, those where a single-gene-knockout results in lethality or severe loss of fitness, have been well studied in many bacterial genomes to develop therapeutic targets for pathogens. Now, stemming from the discovery that the loss of an essential-nearby gene can introduce latent-vulnerabilities specific to cancer cells, the study of human-essential genes has come of practical importance^1^. This importance is magnified as essential genes for cancer-cell growth are found to be located close to target-deletion genes^1^. Therefore, identifying properties of essential genes can further therapeutic developments.

Older genes, with earlier phyletic origin, are more likely to be essential, as well as genes that are hubs in major protein-protein interaction networks^2,3,4^. Essential genes are highly connected with many protein systems, and thus, consistent transcription timing, maintenance of transcript length, and conservation of gene regulation is of high importance^5^. Identification of human essential genes has been approached through the use of single-gene-knockouts, high-throughput mutagenesis, RNAi, and in most recent work, CRISPR–Cas9 editing^6^.

However, moving towards an *in vivo* analysis of gene essentiality, to lend more practical therapeutic insights, studies have focused on the close link between duplication and gene essentiality^7,8^. Duplication is a biological mechanism employed throughout evolution to generate new genetic material^7^. A positive association between singleton, highly-expressed, developmental genes and essentiality is observed, suggesting that essential genes resist duplication events^7,9^. Stemming from these results, copy-number-variants, which result from unequal-crossing-over, retroposition, or chromosomal duplication, were included in efforts to identify essential human genes^1,10^. Intron loss, occurring at an especially greater rate after gene duplication, is the most frequent copy-number-variant in humans, suggesting a likely link between introns and gene-essentiality^11,12^.

Introns, which make-up over half of the non-coding genome, have important regulatory and evolutionary functions. Intron losses and deletions can modulate gene expression patterns and even alter gene function^11^. Typically occurring at the 3’ end of a gene, losses and deletions arise from mediated recombination of a gene with the reverse-transcribed RNA during duplication events or through irregular splice sites^10,13^. Furthermore, intron deletions are most common to longer introns^12^. Intron 1, typically the longest intron, has frequent intron deletions (30.4% of all known deletions) which are especially serious as the first intron preferentially contains regulatory regions and exhibits the highest density of chromatin marks allowing for gene expression^13,14,15^. GC patterns in intronic sequences are associated with an increase in enhancer activity, correct splice site recognition, and protection from intronic deletions^12,16,17^.

It has been suggested that in highly-expressed-genes, selection has resulted in smaller introns that reduce transcriptional cost, which agrees with reports of shorter introns in essential genes^12,18^. Adding to the seeming importance of introns in essential genes, intron deletions in three-essential-yeast genes drastically decreased RNA levels and caused major growth defects^19^.

Owing to the capability of intron losses and deletions to alter gene duplication, expression, and transcription timing, we hypothesize that essential genes, which demand consistency, have developed systems to minimize these events. We thus aim our study to (i) identify whether essential gene introns differ from those of nonessential genes and (ii) characterize the unique properties of essential gene introns to allow for later therapeutic developments.

## Results

We extracted 2135 introns from 165 human essential genes, 74147 introns from 6716 human conditional genes, and 115089 introns from 12449 human nonessential genes from the Ensembl database^20,21^. Human gene essentiality data was gathered from the database of Online Gene Essentiality (OGEE) that gathers experimental data from 18-large-scale-experiments to classify genes by essentiality^3,6^. Conditional genes are genes where experiments have disagreed on essentiality.

We trained a convolutional neural network, based on DeepBind, to predict gene essentiality based on recurring base-pair motifs of 1000 bp long intronic sequence input^22^ (Figure 1). We set aside 20% of the data for the test set and use a three-way random split on the training data to perform three–fold cross– validation for hyperparameter selection. We selected our model’s hyperparameters by performing a grid search of our model’s dropout rate, convolutional layer window size, activation function, and L2 regularization strength. We assessed 36 potential models based on average validation performance across the three folds. The best hyperparameters are then used to train the final model on the entire training set. At training time, we balance our training and validation sets by equally sampling from the essential and nonessential classes. We also ensure that all the introns of a gene lie in the same set so that no gene-specific information affects the validity of our accuracy on the test set. We trained two separate models using the first and last 1000 bp of introns and combined these models by a double classifier which averages essentiality scores from all introns of a gene given by both models. The double classifier optimizes the area under the curve (AUC) of the receiver operating characteristic (ROC) curve used to quantify the diagnostic ability of the model. For the purposes of the neural network, we sought to predict either essentiality or nonessentiality, and thus classified conditional genes from the database as essential if over 50% of experiments agreed on essentiality. If introns of essential genes and nonessential genes have no markedly characteristic properties, we would expect an AUC of 0.5. Rather, our double classifier achieved an AUC of 0.846 (Figure 1).

**Fig. 1:**
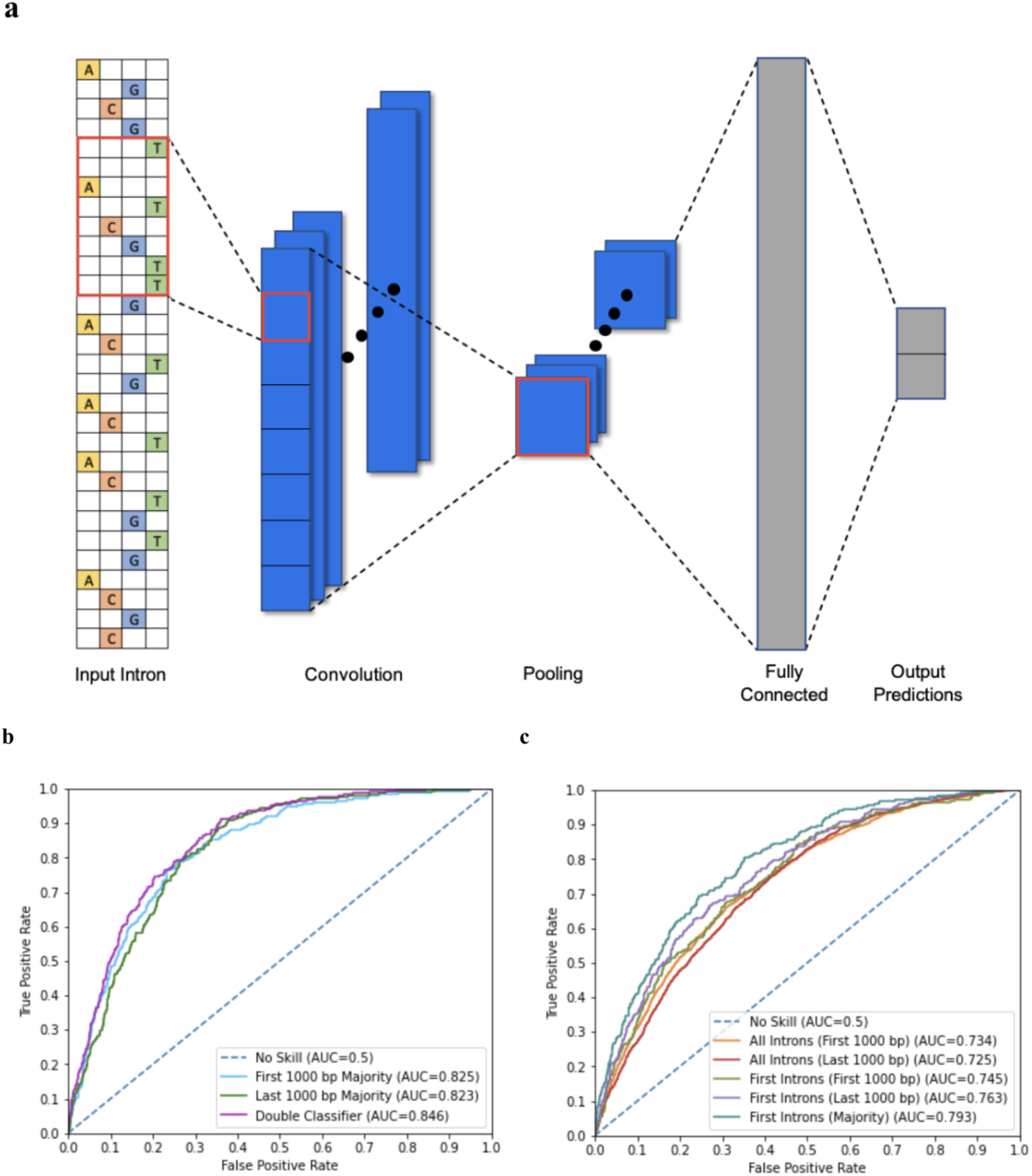
Details of convolutional neural network and testing results. **a**, Our model uses a convolutional architecture to predict intron essentialities. The convolutional layer contains multiple filters that detect motifs within the intronic sequence. Then, the pooling layer averages each filter’s response across the sequence to determine the cumulative presence of motifs. The resulting values are fed into a fully-connected layer followed by a two-value softmax output layer corresponding to the probabilities of the intron being part of an essential or nonessential gene. The best-performing model from our hyperparameter search used 128 convolutional filters with a window size of 24 and a fully connected layer with 128 neurons. We found best results when training with an L2 regularization parameter of 10^−6^ and a dropout rate of 0.2. We trained two models, one on the first 1000 bp of introns and one on the last 1000 bp. This includes the 5’ splice site in the first 1000 bp, as well as the 3’ splice site and the branch site in the last 1000 bp. In all following results, these models are tested on their respective sections of the intronic sequence. **b**, Our model, trained on the first 1000 bp of introns, had an AUC of 0.734. Our model, trained on the last 1000 bp of introns, had an AUC of 0.725. We predicted gene essentiality using a majority classifier on all introns of a gene. The majority classifier of the model trained on the first 1000 bp of introns saw an AUC of 0.825, and the majority classifier of the model trained on the last 1000 bp of introns saw an AUC of 0.823. We further improved accuracy by averaging the outputs of both majority classifiers. This combined classification strategy achieved an AUC of 0.846. **c**, As the first intron is known to have unique properties, we separately tested the models on only first introns, seeing improved accuracy. On first introns, the model trained on the first 1000 bp of introns had an AUC of 0.745 and the model trained on the last 1000 bp of introns had an AUC of 0.763. We further improved first intron essentiality prediction by averaging the outputs of both models to make a dual average prediction, achieving an AUC of 0.793. These results suggest unique properties characterize first introns in essential versus nonessential genes.

Our results demonstrate that introns of essential and nonessential genes have unique properties. To identify these unique properties, we used a computational approach. We also found that the model performs better at classifying introns of strictly essential or nonessential genes, suggesting that conditional genes do not fit well in either essential or nonessential motifs. Therefore, we now include all OGEE classified ‘conditional genes’ as separate entities in our computation to characterize properties of introns by essentiality.

While introns of essential genes differ from introns of nonessential genes by size and number (Figure 2), they also differ by base specific traits. GC motif density is significantly greater in the first introns of essential genes (Figure 3). In order to account for CpG island presence as a potential cofounder of GC density, we show CpG island presence distributions in introns by first or later, and by essentiality. The results of this support that CpG island presence is not a cofounder of GC density results, especially as essential first introns are found to less frequently contain a CpG island than nonessential first introns, and conditional first introns contain the most, which is not parallel to the distribution of GC density among the six classes. Furthermore, we show that GC content, subtracted by GC motif density, is significantly greater in the first introns of essential genes than first introns of nonessential genes as well as later introns of both essential and nonessential genes. However, essential later introns have a significantly lesser GC content than nonessential later introns. GC content has a remarkably similar distribution to GC density (Figure 3).

**Fig. 2:**
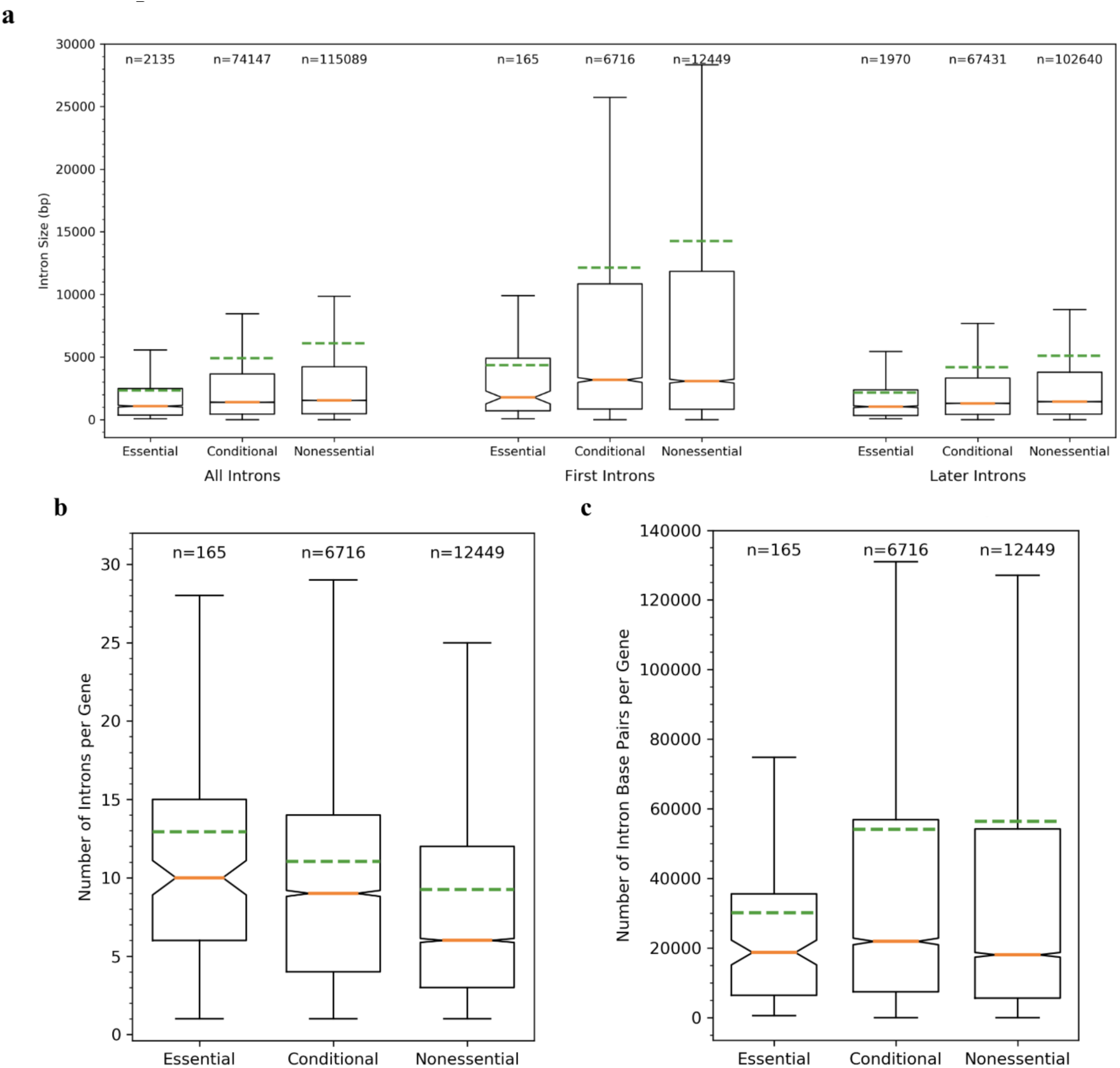
Introns of essential genes differ from introns of nonessential genes by size, number, and position. **a**, The dashed-green line represents the mean and the notches are calculated using a gaussian-based-asymptotic approximation to represent confidence intervals around the medians (orange lines). The first introns for essential (p=0.0001), conditional (p<0.00001), and nonessential (p<0.00001) genes are larger than the gene’s later introns; however, essential gene first introns are longer than the later introns to a lesser degree than those of nonessential introns. The nonessential first intron is much longer (mean times greater) than the essential first intron (p<0.00001). For later introns, nonessential are longer than essential (p<0.00001), but these lengths are closer than the disparity between first intron sizes. Conditional introns typically fall within the middle. **b**, Essential genes have a greater number of introns than both conditional (p=0.021) and nonessential (p<0.00001) genes **c**, However, essential genes have a lesser total length of intronic sequence than both conditional (p<0.00001) and nonessential (p<0.00001) genes.

**Fig. 3:**
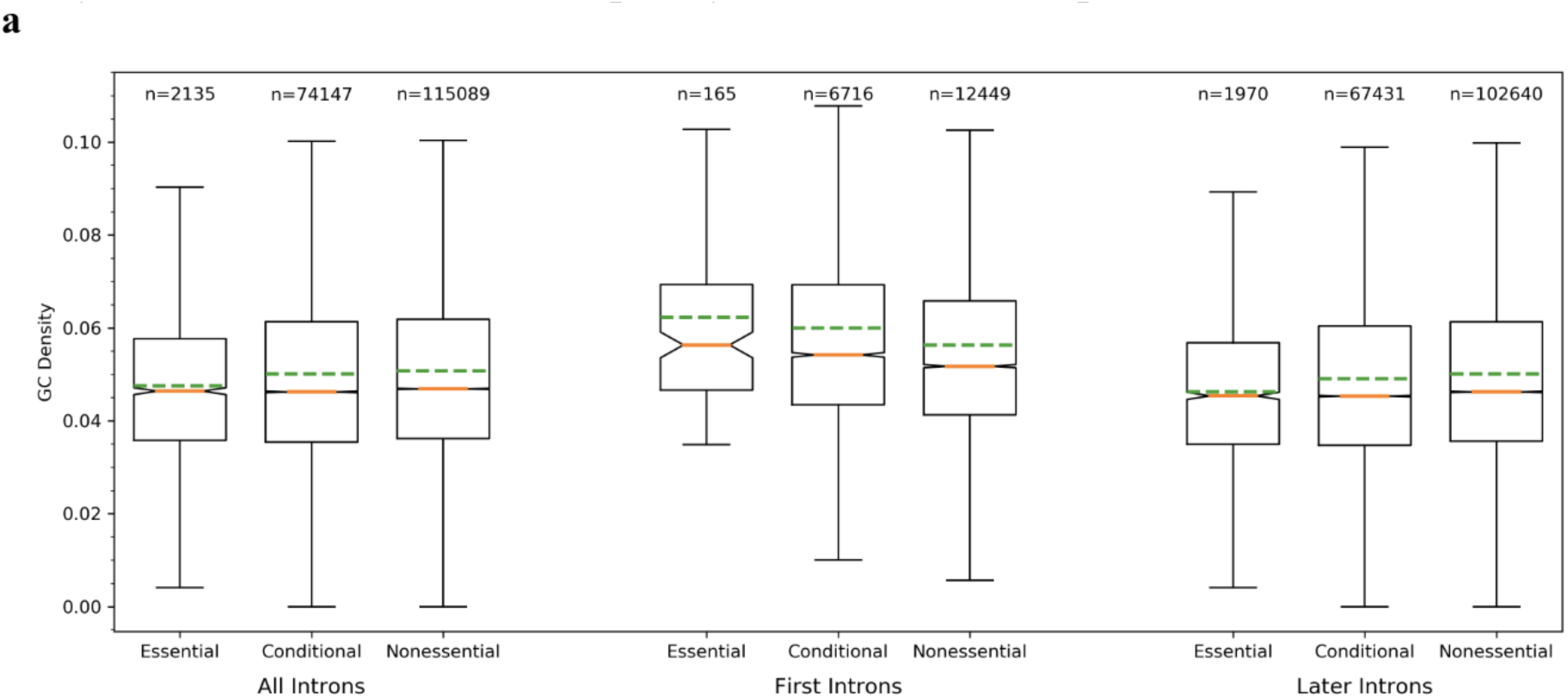
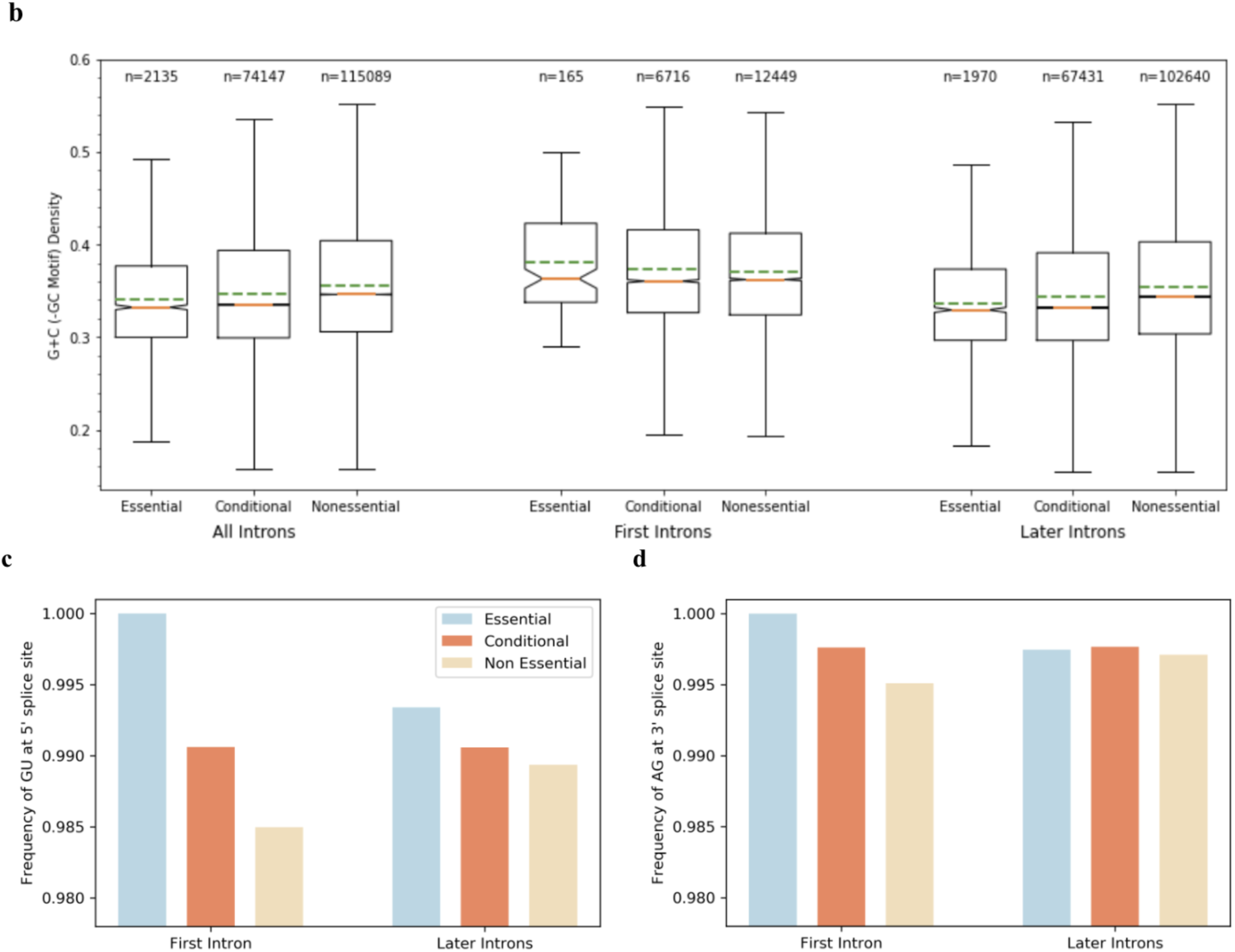
Introns of essential genes differ from introns of nonessential genes by GC motif density, GC content, and lower frequency of unusual 5’ / 3’ splice sites. **a**, The first introns of essential (p<0.00001), conditional (p<0.00001), and nonessential (p<0.00001) genes have a higher GC density than the later introns. Essential (p=0.0004) and conditional (p<0.00001) genes have a higher density of GC regions in their first introns than nonessential first introns. The proportion of GC density of the first intron to later introns for nonessential genes is 1.13, for conditional genes is 1.22, and for essential genes is 1.35. GC density is greater in first introns of essential genes. **b**, GC content, with GC motif content subtracted, has similar distribution to GC motif density among introns split by first/later and essential/conditional/nonessential. GC content is particularly important in annealing strength and increasing gene stability. **c**, Essential gene introns less frequently have an unusual 5’ splice site than conditional introns which in turn less frequently have an unusual 5’ splice site than nonessential introns. The first intron of essential genes is less likely to have an unusual 5’ splice site than conditional or nonessential first introns. Additionally, essential first introns are less likely to have an unusual 5’ splice site than essential later introns. A conditional first intron is less likely to have an unusual 5’ splice site than nonessential first introns, so we see that this effect correlates with essentiality. The first intron of nonessential genes is most likely to have an unusual 5’ splice site. **d**, The first intron of essential genes is less likely to have an unusual 3’ splice site than conditional genes which in turn are less likely to have an unusual 3’ splice site than first introns of nonessential genes. We see that this effect again correlates with essentiality.

Eukaryotic intron 5’ and 3’ splice sites for pre-RNA processing, 5’-GU-AG-3’ boundaries, are highly conserved, while some minor classes of introns have different boundaries^23^. We report that essential gene first introns are less likely to have an unusual 5’ or 3’ splice site when compared to first introns of conditional and nonessential genes. The same trend is true of the 5’ splice site in later introns, to a lesser degree (Figure 3).

Having identified the above features that differentiate introns of essential genes from introns of nonessential genes, we trained a feature–driven deep–learning model to predict gene essentiality so as to determine the importance of the identified features. By only training the model with information on seven features we identified [Average intron size, Number of introns in gene, Intronic bp in gene, GC density (first intron), GC density (later introns), GC count (not including GC motifs) (first intron), GC count (not including GC motifs) (later introns)], we can determine the importance of these identified features in essentiality. Each feature vector corresponds to one gene, where the later intron features correspond to the mean of the value over all later introns. We trained an ensemble deep–learning model to combine results from multiple (10) neural network models so as to reduce variances and generalization errors. The final AUC obtained was 0.787. This provides serious evidence indicating high importance of the seven– identified features in characterizing essential and nonessential genes.

## Discussion

While essentiality is not wholly an intrinsic property of a gene, the ability of our model to predict essentiality or nonessentiality from just intronic sequences, and our later model from just seven features, suggests that there exist characteristic motifs unique to introns of essential genes. The first model’s accuracy for selecting essential introns increases when only testing the first intron as demonstrated by the greater AUC. This suggests that the first introns of essential genes have especially unique characteristics when compared to the first introns of nonessential genes. We followed up on these results with computational analysis of intronic sequences of essential, conditional, and nonessential genes with regard to all introns, only first introns, and only later introns. The primary findings can be summarized in that (i) first introns of essential genes are much shorter than first introns of unessential genes, (ii) essential genes have more introns per gene but these later introns are markedly shorter than the later introns found in nonessential genes, (iii) essential first introns have a greater GC density than first introns of nonessential genes as well as later essential introns, (iv) essential first introns, with essential later introns slightly less so, infrequently have unusual 5’ or 3’ splice sites compared to the first introns of nonessential genes.

Using these features to train feature–based deep learning model to predict gene essentiality, we provide evidence that suggests seven features as major contributing differences between introns of essential and nonessential genes.

From these results, essential genes appear to exhibit intronic characteristics that protect their first introns from loss and deletions. The first intron is crucial for regulation of gene expression; for essential genes which are central to PPI hubs, any deletion in the first intron has the potential to disrupt an entire network^3,14^. Because deletions occur in longer introns at much higher frequency, we found essential first introns are on average over three times less the size of nonessential first introns^12^. First introns of essential genes were found to have a greater GC motif density which allows for an increase in enhancer activity, correct splice site recognition, and protection from intron deletions^12,16,17^. The distribution of GC count closely resembled GC motif density, potentially suggesting that essential first introns have undergone a marked increase in their GC content so as to increase GC motif frequencies for the purposes discussed above. Similarly, as unusual splice sites can allow for alternative splicing, introns of essential genes, especially the first introns, have the lowest frequency of unusual 5’ and 3’ splicing sites^24^. Furthermore, as the majority of deletions occur at the 3’ end, essential genes have an increased number of introns. These later introns however, are smaller than the average nonessential intron, avoiding long introns in essential genes so as to limit any intron loss or deletions. Because deletions in introns of essential genes would alter transcript length and thus interrupt the timing of a complex molecular network, the unique properties of essential introns appear to have been selected to avoid intron losses and deletions.

While we select essential genes based on intronic patterns, other prediction methods of essential genes have been based on support vector machines (SVM), information theoretic statistics, and PPI network leverage^25,26,27^. Information theoretic approaches, trained and tested on the same organism, have shown AUC scores of 0.73 to 0.90 with an average of 0.84, which is slightly lower than our double classifier AUC of 0.846^26^. However, these information–theoretic approaches were not applied to human genes and have lowered accuracy when applied inter-organism. SVM approaches to human essential genes based on 800 selective features report results of mean AUC of 0.8347 and highest AUC of 0.8854, thus with similar accuracy to our intron-based-model and of only slightly increased accuracy to our 7– feature–based deep learning model^28^. PPI–network–leverage combined with feature–information models have shown successes with DeepHe, using PPI network information and 89 sequence derived features, having reported AUC of 0.94 which outperforms SVM, Naive Bayes, Random Forest, and Adaboost models^29^. However, our intron-sequence-model having AUC of 0.846 and our seven–feature–based model having AUC of 0.787 is a surprising result due to the previously unknown role of introns in gene essentiality. With further characterization of introns in essential genes, future feature–based models may rival current PPI approaches.

Conditional genes are correlated between essential and nonessential genes, suggesting a middle ground for both gene stability and alterations of gene functionality. This middle ground is necessary for successful evolution of the genome. We hypothesize that this reflects the desire of the genome to both innovate its genes as well as to conserve its most essential genes. While selecting for deletion–adverse essential intron systems promote basic network stability, while selecting for long, first introns of nonessential genes allows deletions to alter regulation of nonessential genes and even innovate gene function.

The results presented here introduce the concept that essential genes have characteristically unique introns from nonessential genes and we identify 7 features to characterize this difference. These differences, as outlined above, may eventually be exploited to target tumors by disrupting nearby essential genes^1^. Interrupting the complex safety net around the first intron can alter regulation and thereby disrupt a network necessary for tumor growth. Similarly, using targeted CRISPR–Cas9 therapies to force deletions of introns within carefully selected essential genes could likewise stunt cancers.

## Methods

### Model

Our deep learning model is a convolutional neural network based on DeepBind, a predictive model that has shown state–of–the–art performance in predicting sequence specificities of DNA–and– RNA-binding proteins^22^. Our model predicts the essentiality of the gene of an intronic sequence *‘s’* by calculating an essentiality score *f(s) = net(pool(rect(conv(s))))*. Figure 1 depicts our model architecture. Our model accepts 1000 bp sequences encoded as one–hot vectors. The convolutional layer (*conv*) contains multiple filters that detect motifs within the intronic sequence. We apply the ReLU activation function (*rect*), then the pooling layer (*pool*) averages each filter’s response across the sequence to determine the cumulative presence of motifs. The resulting values are fed into a small neural network (*net*) consisting of a fully–connected layer followed by a two–value softmax output layer corresponding to the probabilities of the parent gene being essential or nonessential. The fully–connected layer also uses the ReLU activation function, and the softmax function is applied to the output to normalize prediction probabilities. We prevent our model from overfitting by using L1 and L2 regularization as well as dropout^30^.

### Data

Human DNA sequences and annotations were collected from the Ensembl genome database project^20,21,31^. We used the transcript for each gene corresponding to its longest–coding–sequence as this has been suggested, in recent work, as the most accurate and most biologically relevant^32^. However, both the deep–learning results as well as the bio–computationally–identified feature results were extraordinarily similar when using introns from the longest transcript of each gene. We used the provided annotations to separate out intronic sequences. Before training, the intronic sequences are transformed using one–hot encoding such that each sequence is represented as an *Lx4* matrix for a sequence of length *L*. For our analysis of CpG island presence, we used an algorithm derived by Takai and Jones to detect CpG islands from a nucleotide sequence^33,34^.

We assign labels using gene essentiality information from OGEE, which experimentally classifies genes by essentially^3,6^. OGEE gathers data from 18 databases of large-scale experiments to provide a reference of how many studies found a gene essential or nonessential^3,6^. For the model, due to the ambiguity of conditional genes, we discard all conditional genes that have been found to be essential in less than half of studies. Genes are assigned binary labels of essential or nonessential, where the remaining conditional genes are grouped with essential genes.

We trained two models, one on the first 1000 bp of introns, and one on the last 1000 bp. This includes the 5’ splice site in the first 1000 bp, as well as the 3’ splice site and the branch site in the last 1000 bp. These are the three best characterized regions of eukaryotic introns and are the sites that are most directly involved in spliceosomal modification of the transcript to form mRNA.

### Training Procedure

We set aside 20% of the data for the test set, and use a three–way random split on the training data to perform three–fold cross–validation for hyperparameter selection. We selected our model’s hyperparameters by performing a grid search of our model’s dropout rate, convolutional layer window size, activation function, and L2 regularization strength. We assessed 36 potential models based on average validation performance across the three folds. The best hyperparameters are then used to train the final model on the entire training set. At training time, we balance our training and validation sets by equally sampling from the essential and nonessential classes. We also ensure that all the introns of a gene lie in the same set so that no gene–specific information affects the validity of our accuracy on the test set. We trained all models using Adam gradient descent and a cross–entropy loss minimization objective^35^. The model is trained for 30 epochs with a batch size of 64. We implemented our model using the Keras library running on Tensorflow, and trained on an NVIDIA Tesla M60 GPU.

### Prediction and Evaluation

We evaluate our model on our test set using the area under the curve (AUC) of the receiver operating characteristic (ROC) curve, which measures how well our model distinguishes between essential and nonessential classes. The model produces an essentiality score corresponding to the predicted confidence in the essentiality of the gene of an intron, and the ROC curve is generated by measuring the sensitivity and specificity of the model at varying prediction thresholds of the essentiality score. We also took advantage of both of our models in order to better classify an intron by averaging the scores produced by our two models on the first and last 1000 bp of the intron.

Our model can be extended to classify entire genes with even better accuracy. Rather than classifying the essentiality of individual introns, we classify whether an entire gene is essential or nonessential by combining information from all of its introns. To classify in this manner, we introduce a majority–classification–method. We accept the list of all intronic sequences of a specific gene and run each individual intron through the model to get the essentiality score of each intron. Then we calculate a gene’s essentiality score as the mean of the essentiality scores of its introns.

We attained our highest AUC using a double majority classifier which uses both the first 1000 and last 1000 bp of each intron to classify a gene. We run the first and last 1000 bp from each intron through the models trained on the first and last 1000 bp of each intron, respectively. Then we similarly calculate a gene’s essentiality score as the mean of the essentiality scores of its introns from both models. By combining information from multiple parts of multiple introns, the double–majority–classifier achieves the highest accuracy.

### Feature Model

Our final feature–based model used 7 features extracted from each gene. The normalized feature vectors are used to train a neural network consisting of several fully–connected layers with a binary output. The architecture and hyperparameters of this network were selected by a grid search over the number of hidden layers, number of nodes in each hidden layer, the dropout rate, and L2 normalization strength. Models were evaluated via three–fold cross validation just as with the convolutional model, with the best hyperparameters used to train the final model on the entire test set. We trained all feature models using Adam gradient descent and a cross-entropy loss minimization objective^35^. The model is trained for 30 epochs with a batch size of 64. For final evaluation, we trained an ensemble–deep–learning model to combine results from multiple (10) neural network models so as to reduce variances and generalization errors.

### Code

All the code used for data processing, figure generation, and model training, as well as the weights of our final models, are provided at https://github.com/evendrow/Intron-Essentiality/

## Contributions

Ethan S. and Elan S. conceived the project. E.V. was responsible for data pre-processing. J.V. and E.V. performed the deep–learning and computation. Ethan S., Elan S., J.V., and E.V. analyzed the data. Ethan S. and Elan S. generated conclusions. E.V. and Elan S. created the figures. Ethan S. wrote the manuscript with input from all authors. J.V. and E.V. wrote methods with input from Ethan S.

